# An artificial intelligence-based model for prediction of Clonal Hematopoiesis mutants in cell-free DNA samples

**DOI:** 10.1101/2024.12.11.627785

**Authors:** Gustavo Arango-Argoty, Marzieh Haghighi, Gerald J. Sun, Aleksandra Markovets, J. Carl Barrett, Zhongwu Lai, Etai Jacob

## Abstract

Circulating tumor DNA is a critical biomarker in cancer diagnostics, but its accurate interpretation requires careful consideration of clonal hematopoiesis (CH), which can contribute to variants in cell-free DNA and potentially obscure true tumor-derived signals. Accurate detection of somatic variants of CH origin in plasma samples remains challenging in the absence of matched white blood cells sequencing. Here we present an open-source machine learning framework (MetaCHIP) which classifies variants in cfDNA from plasma-only samples as CH or tumor origin, surpassing state-of-the-art classification rates.

## Main text

Blood-based liquid biopsies are increasingly being explored for personalized cancer diagnostic applications, as a small but detectable amount of circulating tumor DNA (ctDNA) is present in the circulating cell-free DNA (cfDNA) of a blood plasma liquid biopsy^1^. Alongside other signals present in cfDNA^2,3^, noninvasive, potentially longitudinal detection of tumor somatic variants from ctDNA promises to revolutionize early disease detection, prioritization of targeted therapies, identification biomarkers related to favorable response, and tracking of treatment response ^4^.

Isolation of ctDNA variants remains challenging. While removing high allele frequency variants can help filter out germline and somatic non-tumor variants to identify the ctDNA fraction, recent evidence suggests that many low allele frequency variants arise from clonal hematopoiesis (CH) ^5^. CH refers to somatic mutations in hematopoietic cells acquired over a lifetime, often affecting genes commonly altered in hematological malignancies and solid tumors ^6^, such as TP53. CH variants comprise of over 75% of cfDNA variants in noncancerous individuals, and as much as >50% of cfDNA variants in cancerous individuals ^5^. The validity of cfDNA diagnostic applications therefore hinges on accurate detection and removal of CH variants from the putative ctDNA variant pool^7^.

Studies characterizing the presence of CH variants in blood plasma cfDNA commonly utilize a matched nucleated blood cell fraction (e.g. peripheral blood mononuclear cells or white blood cells) to determine variant origin from cfDNA in the absence of a tumor biopsy ^5,8-10^. Such additional samples are not always available in clinical practice. Instead, with only blood plasma sequencing, conventional analyses identify variants as CH by comparing against reference databases of known variants associated with CH or hematological malignancies ^10,11^. Such a knowledge-based approach may suffer from low sensitivity, as many CH variants are known to be specific to individuals or otherwise non-recurrent ^5,8^. The variant allele frequency (VAF) of key hematologic cancer driver genes has also been suggested as a potential metric for distinguishing CH variants in cfDNA samples, although the exact relationship between VAF and the variant’s origin remains unclear^12,13^. Initial efforts have been made to evaluate the feasibility of identifying CH variants using machine learning (ML) -based methods and cfDNA sequencing samples^14^. Although these approaches are still in the nascent stages of research, they highlight the potential for further exploration of ML-based models in the literature.

In this work, we propose an ML-based framework, MetaCHIP, for accurate identification of CH variants from cfDNA samples in the absence of matched sequencing of white blood cells. The proposed framework incorporates layers of information on a variant’s origin derived from trained models on multiple publicly available annotated datasets of blood, tumor and cfDNA samples into a single CH detection framework. The proposed multi-stage framework (summarized in Fig. 1) is fundamentally a meta classifier built on the outputs of two base classifiers. The first base classifier is trained using limited annotated publicly available cfDNA dataset (Fig. 1b). On the other hand, the second base classifier utilizes large publicly available datasets of blood and tumor sequencing samples (Fig. 1c). Each base classifier provides scores on the origin of variants, and a meta classifier is trained to learn the best function for mapping multiple scores to a single final prediction score predicting the variant’s origin (Fig. 1d). All base classifiers use gene and variant embeddings derived from blood and tumor sequencing samples via self-supervised variant representation learning (Fig. 1a, see Methods for details on General-Purpose gene and variant Embeddings). This approach is motivated by the hypothesis that, similar to somatic gene signatures^15^, the sequence context of variants—when combined with prior knowledge of mutations ^16-18^—may reveal information related to the tumor or CH origin of a given variant. Self-supervised learning enables us to capture these representations by leveraging large public datasets of blood and tumor sequencing data and accounting for variant co-occurrences within cancer patients.

**Figure 1:**
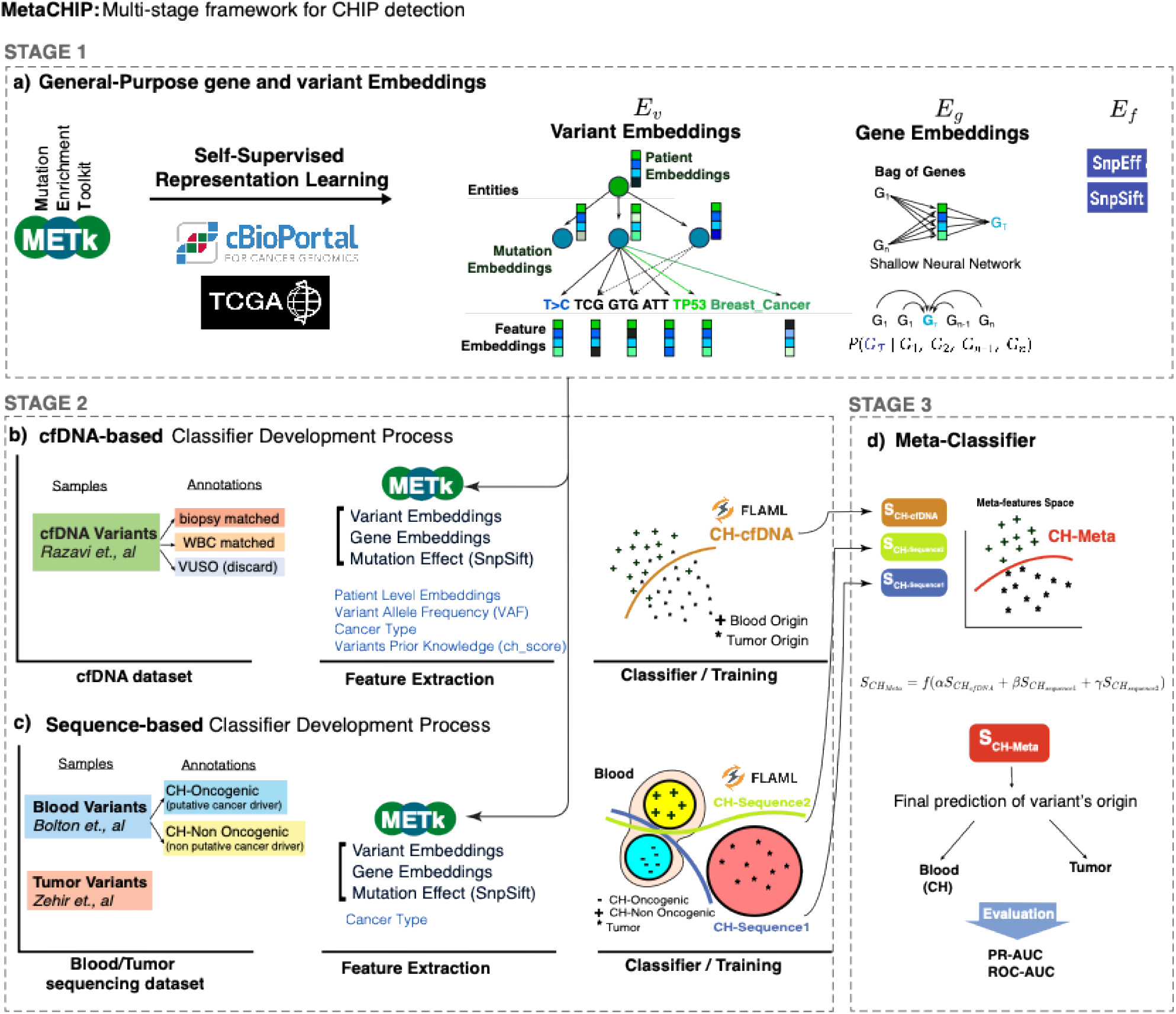
Development of the MetaCHIP framework. (a) Stage 1: variant and gene embedding generation using self-supervised learning. Gene embeddings (*E*g), variant embeddings (*E*v), and functional prediction scores (*E*f) are derived using an Mutation Enrichment Toolkit (METk) , These embeddings are fed to the base classifiers trained in the Stage 2 of the framework. (b) CH-cfDNA classifier. A classifier trained on a cfDNA dataset with experimentally derived annotations (Razavi et al.), designed to score the origin of variants as either CH or tumor. (c) CH-Sequence classifier. A sequence-based classifier trained on large public datasets of tumor and blood genomic samples from MSKCC, scoring variant origin as CH or tumor. CH-Sequence includes two binary classifiers: CH-Sequence1 distinguishes CH-Oncogenic variants from others (tumor or CH-Non Oncogenic), and CH-Sequence2 distinguishes CH-Non Oncogenic variants from others. This model is built on non-cfDNA data to provide independent evidence for variant classification. (d) Meta-classifier (CH-Meta) combining evidence from multiple classifiers. The meta-classifier learn from scores generated by CH-cfDNA and CH-Sequence classifiers to provide a final prediction on variant origin, integrating evidence from cfDNA, tumor, and blood data.

Prior research has demonstrated that larger mutation sequence contexts can enrich information content of genomic data^19^, yet translating these contexts into a compact and meaningful set of numeric features remains an ongoing area of study. More recently researchers have leveraged advances in natural language processing (NLP) to create a shared embedding space for patients and mutations based on the larger-scale sequence context of mutations^20^. All base classifiers in the MetaCHIP framework utilize variant embedding learned from an entity embedding model^21^ that learns representations from the sequence and oncologic context of variants using a self-supervised learning approach (Fig. 1a, described in detail the Methods section). This model generates variant embeddings (*E*_v_), gene embeddings (*E*_*g*_), and functional prediction scores (*E*_*f*_), shown as stage 1 in multi-stage framework. The extracted features in stage 1 are then fed to the base classifiers in stage 2. Together, these features enrich the base classifiers with information of the mutational signature landscape, gene prevalence in cancer, and the functional effect of the variants.

The first base classifier which is the core building block of the MetaCHIP framework, is a cfDNA-based classifier that learns to distinguish the origin of variants in the context of the cfDNA patient’s samples (‘CH-cfDNA, Fig. 1b). The input/output of this classifier is aligned to the input/output of the overall framework in the inference mode. CH-cfDNA model utilizes gene (*E*_g_), variant (*E*_v_) and patient-level embeddings 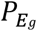 and 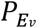, where 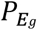 represents the aggregated embeddings of all genes within a patient and 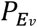 represents the aggregated embeddings of all variants within a patient) from our entity embedding model, as well as sample’s variant allele frequency (*VAF*) and cancer type (Cancer_Type) together with the prior knowledge of CHIP-associated variants (ch_score) and trains a classifier using publicly available cfDNA dataset from *Razavi et al* ^*5*^ which have experimentally derived annotations.

The second base classifier is a sequence-based classifier which takes advantage of large public datasets of patients tumor and blood genomic samples to score variants origin to CH versus tumor (‘CH-Sequence, Fig. 1c).To build this sequence-based classifier, we used two publicly available datasets for CH (blood-derived) and somatic tumor (cancer-derived) mutations from the Memorial Sloan Kettering Cancer Center (MSKCC), which together include 77,068 tumor-derived and 9,810 blood-derived mutations spanning 59 cancer types. Following prior work^22^, we removed duplicate mutations and annotated CH mutations to two subgroups based on their putative role in cancer pathogenesis or their recurrence in myeloid neoplasms to yield a final three class dataset of putative cancer driver (CH-Oncogenic) variants, non-related to cancer pathogenesis (CH-Non Oncogenic) variants, and tumor variants. We next developed an ensemble classifier (‘CH-Sequence’) composed of two binary classifiers that distinguish subcategories of CH variants from the rest of variants. The first, CH-Sequence-1, predicts CH-Oncogenic versus others (tumor or CH-Non Oncogenic; Fig. 1c); the second, CH-Sequence-2: predicts CH-Non Oncogenic versus others (tumor or CH-Oncogenic; Fig. 1c). Note that our CH-Sequence model was trained with data derived from blood and tumor samples and not cfDNA data.

Finally, we apply each of the base classifiers to the cfDNA dataset from *Razavi et al* ^*5*^ to generate scores (*S*_*ch*−*cfDNA*_, *S*_*ch*−*sequence*1_, *S*_*ch*−*sequence*2_) representing the probability of each variant having the blood origin. A meta-classifier (CH-Meta) is trained to learn to predict the origin of variant by integrating the prediction scores from all base classifiers as meta-features (Fig. 1d). We evaluated the standalone performance of each base classifier on the CHIP classification task using cross-validation of the training samples and reported the results as the area under the Precision-Recall (auPR) and the Receiver Operating Characteristic (auROC) curves in Table 1. The cfDNA-based classifier demonstrated superior performance in predicting the variants’ origin, despite being trained on a smaller dataset. This finding highlights the added value of cfDNA samples, which contain patient-level contextual information critical for decoding the origin of variants. The results from the sequence-based classifiers indicate that variants unrelated to cancer pathogenesis (CH-Non-Oncogenic) are significantly more challenging to distinguish from tumor-related or CH-Oncogenic variants. This is expected, as CH-Non-Oncogenic variants are understudied compared to the CH-Oncogenic class and are therefore less prevalent in the datasets used to extract information in stage 1 of the framework. The final row of Table 1 shows the cross-validation performance of the meta-classifier (CH-Meta), revealing a marginal improvement over the cfDNA classifier for predicting variant origins in Razavi’s cfDNA dataset. This suggests that integrating CH-sequence prediction scores as meta-features provides only incremental gains, likely due to the high baseline performance of the cfDNA classifier.

**Table 1:** K-fold (k=5) Cross-validation performance of each framework’s component/classifier.

| **Classifier** | **Stage**<br>**/fig** | **Dataset** | **Annotations/classes**<br>**(N)** | **auROC**<br>**(*mean*±*std*)** | **auPR**<br>**(*mean*±*std*)** |
| --- | --- | --- | --- | --- | --- |
| cfDNA-based<br>(CH-cfDNA) | 2<br>/Fig. b | Razavi | CH vs Tumor<br>(914, 436) | 0.93 ± 0.01 | 0.96 ± 0.01 |
| Sequence-based<br>(CH-Sequence1) | 2<br>/Fig. c | MSK | CH-Oncogenic vs others<br>(2967, 60988) | 0.96 ± 0.003 | 0.78 ± 0.01 |
| Sequence-based<br>(CH-Sequence2) | 2<br>/Fig. c | MSK | CH-Non Oncogenic vs others<br>(3778, 60177) | 0.80 ± 0.01 | 0.27 ± 0.01 |
| Meta-Classifier<br>(CH-Meta) | 3<br>/Fig. d | Razavi | CH vs Tumor<br>(914, 436) | 0.93 ± 0.01 | 0.97 ± 0.01 |

We next evaluated the generalizability of the MetaCHIP framework using four independent external cfDNA validation datasets ^8,23-25^ (summarized in the methods section). A comparison of the predictive power of each classifier on validation datasets consistently showed that CH-Sequence2 was less effective in determining variants origin (Fig. 2a). However, no consistent superiority was observed between CH-Sequence1 and CH-cfDNA, supporting the integration of evidence from both classifiers. CH-Meta, which aggregates all base classifiers prediction scores, demonstrated superior performance compared to individual classifiers across training and most validation datasets (Fig. 2a).

**Figure 2:**
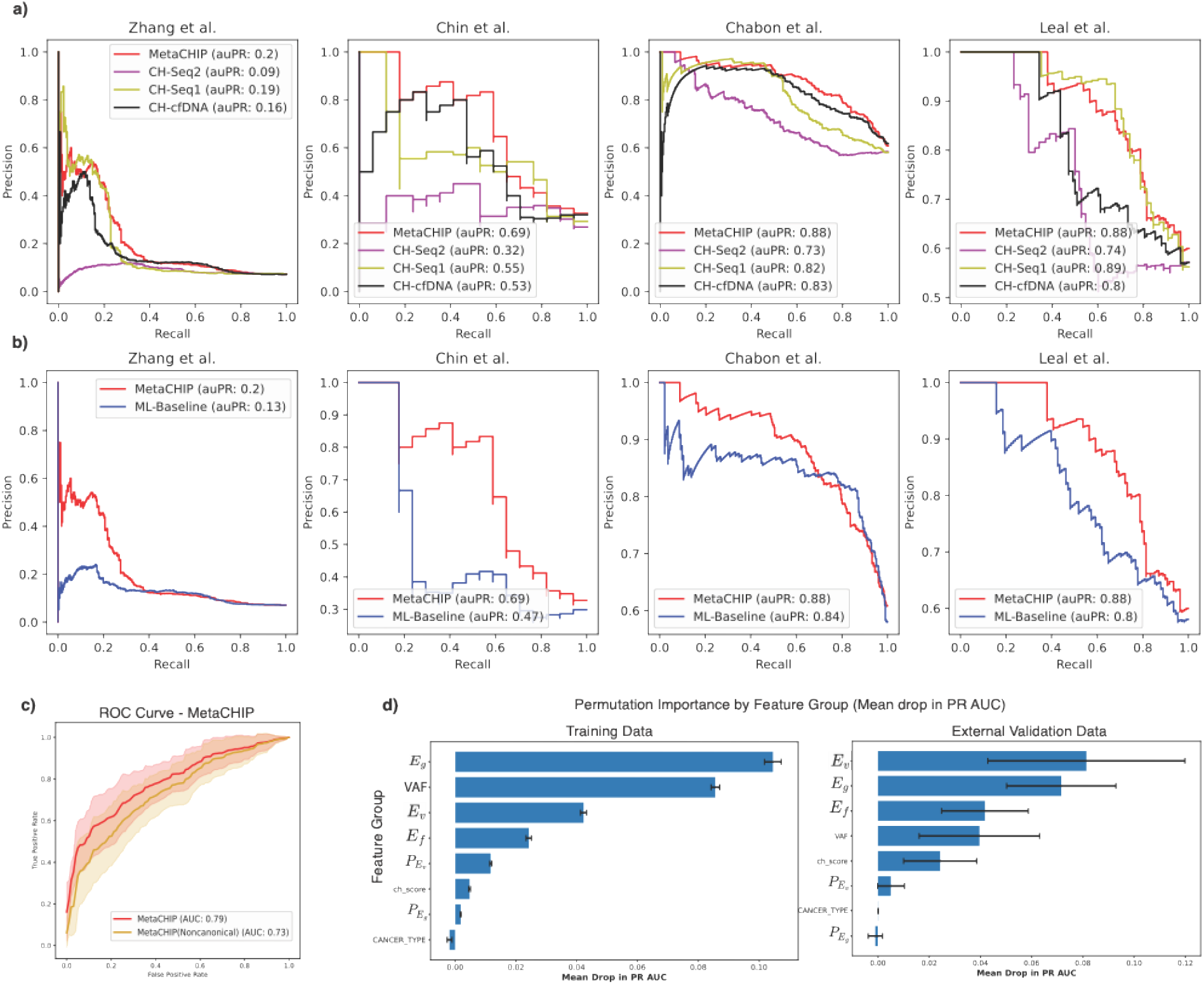
Performance evaluation and benchmarking experiments. a) Precision-recall curves for MetaCHIP and individual classifiers across external cfDNA validation datasets. MetaCHIP outperforms average of the individual classifiers (CH-Seq1, CH-Seq2, and CH-cfDNA) in predictive power for determining variant origin in four independent validation datasets. The area under the precision-recall curve (denoted by auPR) is indicated for each classifier, highlighting MetaCHIP’s consistently higher performance. b) auPR performance comparison of our proposed model (MetaCHIP) versus the machine-learning based baseline model for CHIP classification (ML-baseline) reveals that the proposed model consistently outperforms the ML-Baseline model in identifying the origin of variants across four independent validation datasets. c) Average ROC curves comparing MetaCHIP’s performance with and without canonical CH genes across external validation datasets. MetaCHIP achieves an auROC of 0.79 when canonical CH genes (DNMT3A, TET2, and ASXL1) are included and an auROC of 0.73 when these genes are excluded. ROC curves illustrate overall model performance on the external validation datasets, with shaded areas representing the standard deviation across datasets. d) Permutation feature importance by feature group, measured as mean drop in auPR. Feature importance is assessed across training and external validation datasets, with each bar representing the mean drop in auPR when permuting a specific feature group. In the training data, gene embeddings (*E*g) and VAF emerge as the most influential features. In external validation data, variant embeddings (*E*v) show the highest importance, though with substantial variability across datasets. Gene embeddings (*E*g) exhibit lower variability, indicating their consistent importance across both training and validation datasets. Error bars represent standard deviation across permutation repetitions (n=30) for the training dataset and standard deviation across datasets for the external validation plot.

We benchmarked the MetaCHIP model against an existing machine learning approach for predicting variant origin^14^ (denoted as the *ML-Baseline* model). MetaCHIP consistently outperformed the ML-Baseline model across all external validation datasets (Fig. 2b). Additionally, we examined the effect of removing the most prevalent CH-associated genes (DNMT3A, TET2, and ASXL1), observing a performance drop of approximately 6% in the external validation datasets (Fig. 2c). Notably, MetaCHIP maintained strong performance even when these canonical CH genes were excluded. Finally, we assessed the importance of each feature category in the MetaCHIP model using a permutation importance test (Fig. 2d). Although gene embeddings (*E*_g_) and VAF emerged as the most influential features in the training data, their importance was comparable to variant embeddings (*E*_v_) and functional prediction scores (*E*_f_) in the test datasets. Overall, when considering both training and external validation datasets, gene embeddings (*E*_g_) appear to be the most informative category for determining variant origin (Fig. 2d).

In summary, in this work we have developed a novel modeling framework that predicts CH variants from cfDNA samples with improved performance over the state-of-the-art approaches. This framework achieves this performance level by leveraging gene and variant embeddings derived by self-supervision of large publicly available blood and tumor sequencing datasets and an effective integration strategy to employ them into an ML-based framework for detection of variant’s origin from cfDNA samples. We elaborated the strategy for building and incorporating layers of information on variants’ origin in this framework and demonstrate how each component is contributing to the overall framework’s performance in detecting the origin of the variants. Importantly, we showed that our model generalizes well to independent datasets and can be applied in a pan-cancer fashion.

In the future, CH classification may be further improved with additional patient-level features known to be associated with CH variants, such as age^26^ and prior chemotherapy^22^, as well as other cfDNA-specific features such as fragmentomics^27^ or methylation^28^.

As cfDNA assays and technologies rapidly progress and gain adoption for precision oncology, we believe that our work, combined with existing benchmarking efforts for sequencing ^29^ will enable more robust analyses on critical applications such as early disease detection, minimal residual disease, and mutational profiling to ultimately best inform clinical decision-making.

## Methods

### cfDNA Datasets

Five cfDNA cohorts with reported blood CH and tumor variants were used to train and evaluate the performance of our classifier. The cfDNA dataset from Razavi et al.^5^ includes a prospective study of 124 patients with metastatic cancers, including non-small cell lung, breast, and prostate cancers. This dataset was used to train both the cfDNA-based classifier and the meta-classifier. Razavi et al. performed sequencing on cfDNA, white blood cell DNA, and tumor tissue biopsies for each patient. Notably, 53.2% of the patients harbored mutations indicative of clonal hematopoiesis. The study provides detailed annotation of cfDNA mutations, identifying their origin as blood or tumor. Variants of unknown significance that did not match either blood or tumor were excluded from our analysis. The rest of four additional publicly available cfDNA datasets have been used as external validation datasets ^8,23-25^. *Chabon et al*.^*8*^ describes a personalized deep sequencing method for non-small cell lung cancer (NSCLC) screening. In this study, tumor tissue, pretreatment plasma cfDNA and leukocyte DNA were genotyped in a total of 85 patients with stage I-II-III NSCLC. Similarly, to the *Razavi et al*. study, the majority of cfDNA mutations in this cohort reflect clonal hematopoiesis and are non-recurrent mutations. To validate our classifier using this dataset, only the NSCLC mutations labeled as absent (Tumor) or present (Blood) in matched leukocytes were selected. Chin et al. ^24^ conducted a study focusing on the ultradeep targeted sequencing of circulating tumor DNA (ctDNA) in plasma samples from patients with early and advanced breast cancer. The study included 100 patients, comprising 50 with early-stage and 50 with advanced-stage breast cancer. The researchers employed a targeted sequencing panel covering 1021 cancer-related genes to detect ctDNA mutations. They found that ctDNA was detectable in 43% of early-stage and 86% of advanced-stage breast cancer patients, with higher ctDNA levels correlating with advanced disease stages. The study highlights the potential of ctDNA as a non-invasive biomarker for breast cancer detection and monitoring. Zhang et al.^23^ performed a comprehensive analysis of circulating tumor DNA (ctDNA) across a large cohort of over 10,000 Chinese patients with various cancer types.

The study aimed to assess the prevalence and spectrum of ctDNA mutations in this population. The researchers utilized a targeted sequencing panel to detect ctDNA mutations and found that ctDNA was detectable in 65% of the patients. The mutation profiles varied across different cancer types, with certain mutations being more prevalent in specific cancers. This large-scale study underscores the utility of ctDNA as a biomarker for cancer detection and provides insights into the mutation landscape in the Chinese cancer patient population. Leal et al.^25^ conducted a study focusing on the detection of residual disease in gastric cancer patients through analyses of white blood cells and cell-free DNA (cfDNA). The researchers performed targeted sequencing on cfDNA and matched white blood cell DNA from 50 patients with gastric cancer. They identified mutations in cfDNA that were also present in white blood cells, indicating clonal hematopoiesis (CH). The study highlights the importance of distinguishing between tumor-derived cfDNA mutations and those arising from CH to improve the accuracy of liquid biopsies in detecting residual disease.

### Blood and Tumor Variants Datasets

We used two publicly available datasets for clonal hematopoietic (blood-derived) and somatic tumor (cancer-derived) variants from the Memorial Sloan Kettering Cancer Center (MSKCC). The datasets are available from https://www.cbioportal.org/ under accessions msk_impact_2017 and msk_ch_2020, which together include 77,068 tumor-derived and 9,810 blood-derived mutations spanning 59 cancer types. We used the variant annotation described in Bolton, et al. ^22^ where variants are labeled based on their putative role in cancer pathogenesis using OncoKB ^30^ and its recurrence in a dataset of myeloid neoplasms^31-33^. A total of 5,800 mutations were labeled as Clonal Hematopoietic Putative Cancer Driver (CH-Oncogenic) whereas a total of 4,010 blood mutations were labeled as non-related to cancer pathogenesis (CH-Non Oncogenic). After removing duplicate mutations within each dataset independently, a total of 57,210 tumor mutations, 2,967 CH-Non Oncogenic and 3,778 CH-Oncogenic mutations were used for downstream analysis.

### General-Purpose gene and variant Embeddings: Self-Supervised Representation Learning of genes, variants and patients

Mutational somatic signatures serve as a fingerprint for characterizing the underlying biology of cancer. Tumor somatic mutations tend to leave a strong signature in the cancer cells that can indicate the multiple biological mechanisms present in the tumor (e.g., Aging, Tobacco smoking, APOBEC activity). In cfDNA samples, the enrichment of these signatures may indicate the origin of the mutations ^8,34,35^. As the prevalence of clonal hematopoietic mutations is associated with the aging process, scoring the aging signature for individual mutations can help to identify the origin of blood-derived mutation. In the same context, identifying the tumor mutational signatures may help to identify the mutations that are originated from the tumor. Therefore, identifying the mutational landscape from individual mutations can contribute with the identification of its origin.

Here, we developed a methodology based on neural networks to capture the mutational signature profile of individual variants by learning a shared embedding space for individual mutations, genes, and cancer patients using a self-supervised strategy.

We used the StarSpace^21^ framework to learn a *d*−dimensional space (|*E*_v_|=d) using patient and mutation entities. In detail, the patient entity is described as a bag of all somatic tumor mutations while the mutation entity comprises a set of tokens including the specific mutation, its context, its gene, and cancer type if available (Fig. 1a). During training, one random mutation from a particular patient is selected as the label and the rest of the mutations are selected as input features. At inference time, cancer type is not used as a feature. The mutation context is extracted by taking 2, 3, 4 and 5−mers from the DNA sequence comprising 10 base pairs down and upstream from the mutation’s location are considered as tokens. To train this model we used the pan cancer TCGA dataset along with all available studies in cbioportal (https://portal.gdc.cancer.gov/, downloaded from cbioportal.org). Patient-level variant and gene embeddings (*P*_*E*v_, *P*_*E*g_), were derived by taking the average embeddings for all their variants and gene embeddings respectively (*E*_v,_ *E*_*g*_).

In addition to mutation-based embeddings, we also trained a continuous bag-of-mutated genes model (inspired on the continuous bag-of-words model in natural language processing) to represent mutated genes into a numerical embedding space (*E*_*g*_) (Fig. 1a). We used the pan-cancer mutational dataset from TCGA where each entry is represented by a list of mutated genes. At training time, one gene is randomly selected as label and the remaining are used as features.

Finally, we also extracted a set of functional prediction scores (*E*_*f*_) for non-synonymous mutations using dbNSFP^36^ and SnpSift^16^. These scores are intended to quantify the potential advantage of mutations to carry a positive selection for malignancies. Identifying these mutations is critical for cancer development and can be used to identify cancer-related or clonal hematopoietic mutations.

All these features are available through the <u>M</u>utation <u>E</u>nrichment <u>T</u>ool<u>k</u>it (METk) a python API (to be released upon publication) to automatically extract mutation-related features.

## Incorporating prior knowledge of CHIP-associated variants

DNMT3A, TET2 and ASXL1 are the most prevalent genes in clonal hematopoietic cells ^26^. We extended the list of CH-driver genes by using an in-house list of genes involved in hematopoietic cell regulation, development, and hematological neoplasms. Therefore, any mutation present in any of the 27 CH-related genes are labeled as CH mutations (*U2AF1, BCOR, EZH2, IDH2, MYD88, IDH1, GNAS, STAG2, SF3B1, TET2, MPL, RUNX1, ASXL1, DNMT3A, CREBBP, NPM1, ETV6, BRD4, CBL, JAK2, FLT3, BCL2, STAT3, ZRSR2, CEBPA, SRSF2, PPM1D*). We incorporated this information as a binary feature vector (1 if the mutation is present in CH-driver genes and 0 if it is not)into our framework (denoted as ch_score).

### cfDNA-based Classifier (CH-cfDNA)

cfDNA-based classifier is a binary classifier which classifies each variant in a patient’s cfDNA sample to the blood versus tumor categories (Fig. 1b). This classifier was trained and validated using the cfDNA *Razavi et al*. dataset which consists of 436 tumor somatic mutations and 914 blood CH derived mutations. Features used to train this classifier were variant and gene embedding (*E*_v_, *E*_*g*_), patient-level embedding (*P*_*E*v_, *P*_*E*g_), and functional prediction scores (*E*_*f*_) derived by METK and variant per patient-level *VAF* and cancer type.

### Sequence-based Classifier (CH-Sequence)

The CH-Sequence model is not trained on data obtained directly from cfDNA datasets, rather it uses data derived from blood and tumor samples. This strategy has the advantage of using evidence derived from a classifier that can leverages sequence-based cancer-context aware representations at the mutation level extracted from large training datasets, to contribute extra evidence on the origin of variants to the final CHIP detection framework.

To predict the origin of mutations (blood vs tumor) we built two specialized classifiers. One classifier (CH-Sequence1) to detect blood CH-Oncogenic from other mutations and a second classifier (CH-Sequence2) to detect blood CH-NO from other mutations (Fig. 1c). The CH-Sequence1 classifier was trained using the CH-Oncogenic mutations as the primary class while the CH-NO and tumor mutations were used as the secondary class. For the CH-Sequence2 classifier CH-NO mutations were labeled as the primary class while CH-Oncogenic and tumor mutations were labeled as the secondary class (Fig. 1c). Then, each mutation has three predictive (CH-NO, CH-Oncogenic and tumor) scores. The tumor origin score is defined as the mean value between CH-Sequence1 and CH-Sequence2 tumor scores and the blood origin score is defined as the mean between CH-NO and CH-Oncogenic scores.

The set of features used to train the sequence-based classifiers are the mutation embeddings (|*E*_*m*_|=128), the gene level embeddings (|*E*_*g*_|=8), the functional prediction scores (|*E*_*f*_|=37) and the cancer type (|*C*_*t*_|=1).

### Meta Classifier (CH-Meta)

In order to combine the advantages of large non-cfDNA datasets (used to train sequence-based classifiers) and cfDNA-specific features (used to train cfDNA classifier), we trained a meta classifier (CH-Meta) which takes the predicted probability scores for the CH class according to each of the base classifiers as the input (meta-features) and learns a function to distribute weights across various sources of evidence 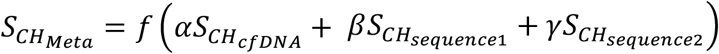. We used a logistic regression classifier to learn this function. To avoid overfitting, we used the same training and validation dataset splits employed in training the cfDNA-based classifier, as both classifiers were trained on the same data samples but with different features.

### Model Selection and Performance Evaluation

We conducted stratified 5-fold cross-validation to evaluate the performance of each classifier contributing evidence to the final model. Both the area under the Receiver Operating Characteristic curve (auROC) and the area under the Precision-Recall curve (auPR) were used as evaluation metrics. Due to the class imbalance in most datasets, we primarily focused on auPR for result interpretation, while auROC is reported where relevant. Each evidence-generating classifier was trained using FlaML (Fast and Lightweight AutoML), a Python library that automates model selection and efficiently tunes hyperparameters^37^.

## References

1 Diaz Jr, L. A. & Bardelli, A. Liquid biopsies: Genotyping circulating tumor DNA. Journal of Clinical Oncology 32, 579–586 (2014). 10.1200/JCO.2012.45.2011

2 Snyder, M. W., Kircher, M., Hill, A. J., Daza, R. M. & Shendure, J. Cell-free DNA Comprises an In Vivo Nucleosome Footprint that Informs Its Tissues-Of-Origin. Cell 164, 57–68 (2016). 10.1016/j.cell.2015.11.050

3 Liu, M. C. et al. Sensitive and specific multi-cancer detection and localization using methylation signatures in cell-free DNA. Ann Oncol 31, 745–759 (2020). 10.1016/j.annonc.2020.02.011

4 Alix-Panabières, C. & Pantel, K. Clinical applications of circulating tumor cells and circulating tumor DNA as liquid biopsy. Cancer Discovery 6, 479–491 (2016). 10.1158/2159-8290.CD-15-1483

5 Razavi, P. et al. High-intensity sequencing reveals the sources of plasma circulating cell-free DNA variants. Nature medicine 25, 1928–1937 (2019).

6 Steensma, D. P. et al. Clonal hematopoiesis of indeterminate potential and its distinction from myelodysplastic syndromes. Blood 126, 9–16 (2015). 10.1182/blood-2015-03-631747

7 Hu, Y. et al. False-Positive Plasma Genotyping Due to Clonal Hematopoiesis. Clin Cancer Res 24, 4437–4443 (2018). 10.1158/1078-0432.CCR-18-0143

8 Chabon, J. J. et al. Integrating genomic features for non-invasive early lung cancer detection. Nature 580, 245–251 (2020). 10.1038/s41586-020-2140-0

9 Jensen, K. et al. Association of Clonal Hematopoiesis in DNA Repair Genes With Prostate Cancer Plasma Cell-free DNA Testing Interference. JAMA Oncol 7, 107–110 (2021). 10.1001/jamaoncol.2020.5161

10 Yaung, S. J. et al. Clonal Hematopoiesis in Late-Stage Non–Small-Cell Lung Cancer and Its Impact on Targeted Panel Next-Generation Sequencing. 1271–1279 (2020). 10.1200/po.20.00046

11 Si, H. et al. A Blood-based Assay for Assessment of Tumor Mutational Burden in First-line Metastatic NSCLC Treatment: Results from the MYSTIC Study. 27, 1631–1640 (2021). 10.1158/1078-0432.CCR-20-3771 %J Clinical CancerResearch

12 Adebayo, J. et al. Sanity checks for saliency maps. Advances in neural information processing systems 31 (2018).

13 Young, A. L., Tong, R. S., Birmann, B. M. & Druley, T. E. Clonal hematopoiesis and risk of acute myeloid leukemia. haematologica 104, 2410 (2019).

14 Fairchild, L. et al. Clonal hematopoiesis detection in patients with cancer using cell-free DNA sequencing. Science translational medicine 15, eabm8729 (2023).

15 Tate, J. G. et al. COSMIC: the Catalogue Of Somatic Mutations In Cancer. Nucleic Acids Res 47, D941–D947 (2019). 10.1093/nar/gky1015

16 Cingolani, P. et al. Using Drosophila melanogaster as a Model for Genotoxic Chemical Mutational Studies with a New Program, SnpSift. Front Genet 3, 35 (2012). 10.3389/fgene.2012.00035

17 Cingolani, P. et al. A program for annotating and predicting the effects of single nucleotide polymorphisms, SnpEff: SNPs in the genome of Drosophila melanogaster strain w1118; iso-2; iso-3. Fly (Austin) 6, 80–92 (2012). 10.4161/fly.19695

18 Landrum, M. J. et al. ClinVar: improving access to variant interpretations and supporting evidence. Nucleic Acids Res 46, D1062–D1067 (2018). 10.1093/nar/gkx1153

19 Stobbe, M. D. et al. Recurrent somatic mutations reveal new insights into consequences of mutagenic processes in cancer. PLoS computational biology 15, e1007496 (2019).

20 Zhang, Y., Xiao, Y., Yang, M. & Ma, J. Cancer mutational signatures representation by large-scale context embedding. Bioinformatics 36, i309–i316 (2020).

21 Wu, L. et al. StarSpace: Embed All The Things! , 1709.03856 (2017). <https://ui.adsabs.harvard.edu/abs/2017arXiv170903856W>.

22 Bolton, K. L. et al. Cancer therapy shapes the fitness landscape of clonal hematopoiesis. Nat Genet 52, 1219–1226 (2020). 10.1038/s41588-020-00710-0

23 Zhang, Y. et al. Pan-cancer circulating tumor DNA detection in over 10,000 Chinese patients. Nat Commun 12, 11 (2021). 10.1038/s41467-020-20162-8

24 Chin, Y. M. et al. Ultradeep targeted sequencing of circulating tumor DNA in plasma of early and advanced breast cancer. Cancer Science 112, 454–464 (2021).

25 Leal, A. et al. White blood cell and cell-free DNA analyses for detection of residual disease in gastric cancer. Nature communications 11, 525 (2020).

26 Bick, A. G. et al. Inherited causes of clonal haematopoiesis in 97,691 whole genomes. Nature 586, 763–768 (2020). 10.1038/s41586-020-2819-2

27 Cristiano, S. et al. Genome-wide cell-free DNA fragmentation in patients with cancer. Nature 570, 385–389 (2019). 10.1038/s41586-019-1272-6

28 Shen, S. Y. et al. Sensitive tumour detection and classification using plasma cell-free DNA methylomes. Nature 563, 579–583 (2018). 10.1038/s41586-018-0703-0

29 Deveson, I. W. et al. Evaluating the analytical validity of circulating tumor DNA sequencing assays for precision oncology. Nat Biotechnol 39, 1115–1128 (2021). 10.1038/s41587-021-00857-z

30 Chakravarty, D. et al. OncoKB: A Precision Oncology Knowledge Base. JCO Precis Oncol 2017 (2017). 10.1200/PO.17.00011

31 Grinfeld, J. et al. Classification and Personalized Prognosis in Myeloproliferative Neoplasms. N Engl J Med 379, 1416–1430 (2018). 10.1056/NEJMoa1716614

32 Papaemmanuil, E. et al. Somatic SF3B1 mutation in myelodysplasia with ring sideroblasts. N Engl J Med 365, 1384–1395 (2011). 10.1056/NEJMoa1103283

33 Papaemmanuil, E. et al. Genomic Classification and Prognosis in Acute Myeloid Leukemia. N Engl J Med 374, 2209–2221 (2016). 10.1056/NEJMoa1516192

34 Cook, J. H., Melloni, G. E. M., Gulhan, D. C., Park, P. J. & Haigis, K. M. The origins and genetic interactions of KRAS mutations are allele- and tissue-specific. Nat Commun 12, 1808 (2021). 10.1038/s41467-021-22125-z

35 Alexandrov, L. B. et al. The repertoire of mutational signatures in human cancer. Nature 578, 94–101 (2020). 10.1038/s41586-020-1943-3

36 Liu, X., Li, C., Mou, C., Dong, Y. & Tu, Y. dbNSFP v4: a comprehensive database of transcript-specific functional predictions and annotations for human nonsynonymous and splice-site SNVs. Genome Med 12, 103 (2020). 10.1186/s13073-020-00803-9

37 Wang, C., Wu, Q., Weimer, M. & Zhu, E. FLAML: A Fast and Lightweight AutoML Library. Fourth Conference on Machine Learning and Systems (MLSys 2021) (2021).

